# Molecular drivers of mutualistic association between anemone and anemonefish

**DOI:** 10.1101/2025.08.21.671668

**Authors:** Sneha Suresh, Daniele Romeo, Celia Schunter

## Abstract

The anemone-anemonefish mutualism is one of the most iconic in the marine environment. While the evolution of this mutualistic relationship has contributed to the ecological success of both partners, the underlying molecular processes that establish and maintain it remain poorly understood, particularly how anemonefish tolerate anemone venom. Here we explored the transcriptional dynamics in the anemonefish *Amphiprion clarkii* and its host anemone *Entacmaea quadricolor* 48 hours after association to identify the molecular underpinnings of their mutualistic relationship. Upon acclimation with an anemone, anemonefish showed differential regulation of sensory perception and memory genes in key brain regions, indicating activation of neural pathways that may facilitate host recognition and mutualism establishment. In the fish’s skin, altered expression of genes involved in neurotransmitter release, cytoskeleton organization, and venom receptor proteins points to mechanisms of resistance to anemone venom. This resistance is particularly remarkable since anemone hosting fish exhibited increased expression of genes encoding mechanoreceptors, venom proteins, and ion channels involved in nematocyst discharge, indicating the anemone does indeed mount an active response to their mutualistic partner. Overall, our results highlight the complex interplay of molecular events in both species that play a pivotal role in establishing this mutualistic relationship.

## Introduction

Mutualistic interactions are major drivers of biodiversity and ecosystem function, influencing evolutionary trajectories across diverse taxa [1]. These complex associations evolve through various processes [2,3] and, once established, can drive evolutionary transitions and species diversification [2,4]. For instance, mutualisms between plants and mycorrhizal fungi, plants and pollinators, corals and dinoflagellates, sea anemones and fish, as well as primates and plants, have all contributed to evolutionary diversifications and increased species richness [3,5,6].

Among marine mutualisms, the association between sea anemones and anemonefishes is especially intriguing. Sea anemones produce venom both in their mucous secretions and through nematocysts discharged upon mechanical or chemical stimulation as a defence mechanism [7]. While the venom produced by sea anemones is lethal to most marine organisms, anemonefishes exhibit a unique resistance enabling them to form mutualistic relationships with one or more of ten sea anemone species [8,9]. This mutualistic association has enhanced the ecological success of both partners [10]. Anemonefish gain protection from predation and a secure breeding site [10], while the host anemones benefit from defense against predators, nutrient input, and increased oxygenation that promote growth and reproduction [11]. Notably, while anemonefishes are obligate symbionts and never found solitary in nature, anemones can survive independently [9]. This obligate association was contingent on anemonefishes developing specific molecular and physiological adaptations to live unharmed among the anemone’s stinging tentacles [12]. Several hypotheses have been proposed to explain this resistance, with the anemonefish’s mucous coating widely considered to play a key role in protecting them from anemone toxins [13–15]. The mucus layer of anemonefishes is proposed to be thicker, chemically distinct from other coral reef fishes, and lacking compounds that trigger nematocyst discharge [11,15]. In particular, anemonefish mucus contains lower levels of sialic acids, a compound that triggers nematocyst firing [13,16], and may also be molecularly similar to anemone mucus, effectively acting as a “chemical camouflage” to inhibit nematocyst discharge [17,18].

A key aspect of this resistance also involves a behavioural process called “acclimation” during initial contact with a host anemone. During acclimation, fish perform stereotyped behaviours in which they repeatedly touch the anemone tentacles first with only their fins and then gradually increasing body contact with the tentacles until they are no longer stung [17]. During this acclimation process, molecular changes likely occur within the fish that confer venom resistance [11]. Recent genomics studies identified genes that underwent positive selection during anemonefish diversification, such as Versican core protein and Protein O-GlcNAse, which alters the structure of N-acetylated sugars in fish mucus glycoproteins thereby preventing nematocyst discharge and contributing to anemonefish venom resistance [12]. However, molecular changes occurring in the anemonefish during the critical acclimation period remain uncharacterized, limiting our understanding of the mechanisms of venom resistance.

While most research has focused on the fish partner, the host anemone may also undergo physiological and molecular changes during acclimation. Recent studies have shown that the microbiomes of both the host anemone and anemonefish are altered after association [19–22]. Furthermore, if acclimated anemonefish no longer triggers nematocyst discharge, this should be reflected in the host anemone through reduced venom production upon acclimation and mutualistic association. Therefore, investigating how venom production changes upon mutualism establishment can provide crucial insights into the molecular mechanisms underlying the formation and maintenance of this complex mutualistic relationship.

The present study investigates the molecular underpinnings of mutualism between the sea anemone *Entacmaea quadricolor* and the anemonefish *Amphiprion clarkii*, both of which are generalist species [8,23,24]. *E. quadricolor* is a natural host of *A. clarkii* [11,25] and *A. clarkii* requires an acclimation period in order to fully associate with *E. quadricolor* [21,25,26]. Additionally, when separated from its host anemone for prolonged durations, *A. clarkii*, similar to other anemonefish, needs to reacclimate [18,27]. This acclimation period plays a crucial role in establishing the symbiotic relationship, and we hypothesize that during acclimation gene expression patterns are altered in both partners. By investigating the transcriptional landscape of both species upon acclimation, we aim to identify (1) the molecular changes in both the host anemone and the anemonefish that facilitate mutualistic association and (2) the molecular processes conferring resistance of the anemonefish to the anemone venom.

## Materials and Methods

### Sample collection and experimental design

*Amphiprion clarkii* and *Entacmaea quadricolor* were chosen as model species for our study as both are widely distributed and *E. quadricolor* is the most common host anemone for *A. clarkii* in Hong Kong [28]. Adult *A. clarkii* individuals were collected by SCUBA diving from a depth of 5-7 meters using barrier nets and hand nets in September 2021 from two locations (N 22°15.525’ E 114°20.981’ and N 22°15.545’ E 114°20.930’) in South Ninepin Island, Hong Kong. Fish were transported to the aquarium facility at The University of Hong Kong and maintained for a period of seven months (until April 2022) in complete isolation from anemones, which is sufficient time for anemonefish to lose their protection from anemone venom [11,27].

Ten individuals of *E. quadricolor*, originally collected in Hong Kong waters, were purchased from a local aquarium store and maintained in isolation from anemonefish in separate tanks for a period of five days prior to the start of the experiment. A shorter isolation time was chosen for anemones as it has been previously established that re-acclimation of fish with anemones is not influenced by the length of time anemones have been in isolation [27]. Both fish and anemones were maintained in 30L tanks with a photoperiod of 12/12hr (light/dark cycle). Fish and anemones were fed *ad libitum* with commercial feed once daily, with fish diet supplemented with brine shrimp once per week. Temperature, pH, dissolved oxygen, and salinity were kept constant throughout the experiment while mimicking the natural parameters of the collection site (Supplementary Table S1).

At the start of the experiment, five anemones were transferred to five different tanks, each containing one fish. For the control treatment, four artificial anemones (made from rubber bands) were transferred to four other tanks, each containing one fish. Five control anemones (kept in isolation throughout the experiment) were also transferred between tanks to account for movement effects (Figure 1). The interaction between fish and anemones was recorded for five hours per day for two consecutive days using SQ23 Wireless HD 1080P mini cameras to observe the acclimation behaviour and ensure that acclimation was complete prior to tissue sampling. Acclimation was considered complete when fish could swim freely among anemone tentacles without any visible distress or avoidance. No water changes were conducted during the experiment, and the water-flow was turned off to avoid altering olfactory cues that anemonefish rely on to recognize anemones. Fish and anemones were allowed to interact for a period of 48 hrs, a duration shown to be sufficient for full acclimation [11,27], which was also confirmed by observations in our study. A similar time frame of interaction was used in a previous study investigating whether resistance of fish to anemones is facilitated by transfer of mucous antigens from anemones to fish [17]. After 48 hrs, all fish were euthanized by severing the spine, standard length was measured, and skin and five brain regions (diencephalon, telencephalon, optic tectum, cerebellum, and brain stem) were dissected and snap frozen in liquid nitrogen. All fish were sexed by visual examination of their gonads. A total of ten tentacles were sampled from each individual anemone at the end of the experiment and snap frozen in liquid nitrogen. The pedal disc diameter of all anemones was measured prior to dissection. All tissue samples were stored at - 80°C until further processing.

**Figure 1:**
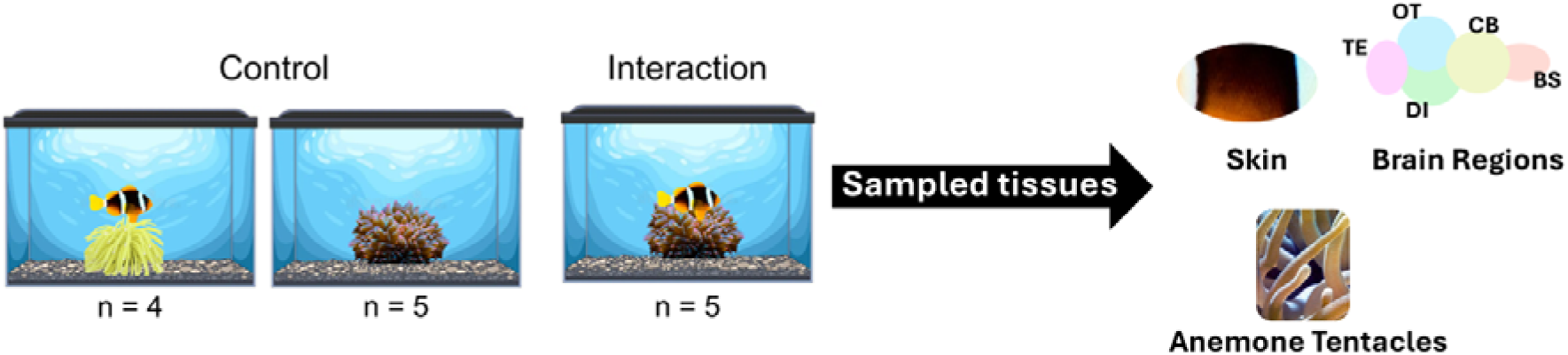
Schematic representation of the experimental design showing from left to right: control fish (with fake anemone), control anemone and physical interaction between the fish and anemone and the sampled tissues for RNA sequencing. BS: Brain stem, CB: Cerebellum, DI: Diencephalon, OT: Optic tectum, TE: Telencephalon.

### RNA extraction, sequencing, and sequence processing

Frozen skin tissue samples from *A. clarkii* were homogenised in buffer RLN (Qiagen), that was pre-cooled to 4°C, for 5 mins at 30 cycles/s using Qiagen TissueLyzer II with 10-15 single-use silicon beads per sample. Frozen brain regions were homogenised in buffer RLN (Qiagen), that was precooled to 4°C, using sterile micropestles. Homogenates were then centrifuged, and the supernatant was mixed with buffer RTL. The Qiagen RNeasy Mini Kit was subsequently used to extract total RNA. On-column DNase I treatment (Qiagen) was incorporated during the extraction process, according to the manufacturer’s instructions, to remove potential DNA contamination.

Total RNA was extracted from frozen anemone tentacles following the protocol from [29], with slight modifications using Phasemaker tubes (Invitrogen). Frozen tissue samples were directly homogenised in TRIzol™ (Life Technologies) using sterile micropestles, and a second extraction step was performed by adding 250 µl of TRIzol™ and 250 µl of Chloroform:isoamyl alcohol (24:1; Sigma-Aldrich). To remove any DNA contamination, in-solution DNase digestion was performed using DNAse 1 (Qiagen) followed by RNA cleanup (Qiagen) according to the manufacturer’s instructions.

The concentration of extracted RNA from all the samples was measured using Qubit and RNA quality was determined using Agilent 2100 Bioanalyzer. Samples with an RNA Integrity Number (RIN) > 6.5 were used for cDNA library preparation using KAPA mRNA HyperPrep Kit with one hundred nanograms of total RNA as starting material. The mRNA samples were subsequently sequenced on Illumina NovaSeq 6000 to generate 151bp paired-end reads at the Centre for PanorOmic Sciences, The University of Hong Kong.

Quality of all raw reads was determined using FastQC [30] v0.11.8. Adapters and low quality reads (Phred scaled quality score < 20 for fish skin and brain and < 25 for anemone) were removed using Trimmomatic [31] v0.39 with a 4-base sliding window approach using the parameters ILLUMINACLIP:adapters.fa:2:30:15:8:true MINLEN:40. Different quality thresholds were chosen for anemone and fish samples due to differences in average quality of raw reads. Additionally, for anemone samples, over-represented sequences reported by FastQC were also removed. The trimmed sequences from both species were further filtered to remove potential contaminant sequences from bacteria, fungus and virus using Kraken [32] v2.0.8-beta with a confidence score of 0.7 and 0.3 for *A. clarkii* and *E. quadricolor*, respectively.

An average of 24 ± 4.1 and 31 ± 2.2 million high quality reads from *A. clarkii* skin and brain tissues (Supplementary Table S2) were then mapped to the reference genome [33] using HISAT2 [34] v2.2.1 (parameters --no-mixed --no-discordant). On average, 70.68 ± 12.61% and 83.68 ± 4.23% of the sequenced reads from skin and brain samples, respectively, mapped to the reference genome (Supplementary Table S2). FeatureCounts [35] v2.0.6 was subsequently used to obtain raw read counts per gene with multi-mapped reads being assigned fractional counts (parameters: -M --fraction -B -p --countReadPairs).

### *De novo* transcriptome assembly and annotation for *E. quadricolor*

Currently, no reference genome exists for *E. quadricolor*, so a *de novo* transcriptome assembly was constructed. Adapter-free, quality trimmed and decontaminated sequences of *E. quadricolor* were further filtered using the ‘filter_illumina’ script from DRAP [36] v1.92 to identify and remove reads containing ambiguous bases (Ns). Specifically, for all reads containing ambiguous bases, the longest subsequence without ‘N’ was extracted and retained if it constituted at least half of the original sequence length and was 40bp or longer. An average of 23 ± 1.5 million read pairs from eight individuals (which retained the most reads after all filtering steps; Supplementary Table S2) were merged and assembled *de novo* using DRAP with Trinity [37] as the assembler using the pooled assembly strategy (parameters: -c ’-x’ --no-trim --no-norm -m bwa --no-rate). The initial transcriptome assembly consisted of 463,812 contigs.

Since sea anemones contain endosymbiotic dinoflagellates (Symbiodiniaceae), the initial transcriptome assembly contained a mixture of anemone and dinoflagellate transcripts, as evidenced by two peaks in the GC content plot (Supplementary Figure S1(a)). Anthozoans and dinoflagellates have drastically different GC contents, with Anthozoa having ∼40% and Symbiodiniaceae having ∼55% [38]. This difference in GC content can be used to separate transcripts belonging to dinoflagellates from those of the anemone as done previously [38]. To disentangle the meta-transcriptome, the initial transcriptome assembly was blasted (using BLASTN; parameters: -evalue 1e-20 -max_hsps 1) against a collection of Symbiodiniaceae genomes obtained from the NCBI [39] and SAGER [40] databases (Supplementary Table S3). All transcripts with BLAST hits to Symbiodiniaceae and GC content above 40% were removed from the transcriptome assembly. The resulting “cleaned” transcriptome consisted of 397,169 contigs and only a single peak in the GC content plot (Supplementary Figure S1(b)). Of the 66,643 transcripts identified as dinoflagellate sequences and removed from the initial *de novo* assembly, the majority were from the genus *Cladocopium* (Supplementary Figure S1(c)). This is similar to the findings of [38], who found that sea anemones from the fringing reefs of Okinawa form symbiotic relationships predominately with *Cladocopium*.

To reduce redundancy in the transcriptome assembly, TransDecoder [41] v5.5.0 was used on the “cleaned” assembly to identify transcripts containing coding regions (ORFs), and only those containing ORFs over 100 amino acids were retained. These were then blasted against the UniProtKB/Swiss-Prot Release 2023_01 database (-evalue 1e-5 -max_target_seqs 1) and only the best ORF from each contig was retained, prioritized first based on blast homology and then based on ORF length. A final clustering step was performed using Corset [42] v1.09, which hierarchically clusters contigs to genes, and the longest transcript per cluster was extracted using the fetchClusterSeqs.py script from Corset-tools (https://github.com/Adamtaranto/Corset-tools). The final transcriptome assembly consisted of 38,887 high-quality contigs. Completeness of the final transcriptome assembly was determined using BUSCO v4.1.3 [43] using the Metazoa_odb10 set of Benchmarking Universal Single-Copy Orthologs. The percentage of complete BUSCOs was above 90% with only a small fraction (6.4%) being duplicated (C:91.5% [S:85.1%, D:6.4%]; Supplementary Figure S2), indicating that the assembly is highly complete. Basic statistics of the transcriptome assembly was determined using ‘TrinityStats.pl’ from Trinity accessory scripts (https://github.com/trinityrnaseq/trinityrnaseq/tree/master/util), rnaQUAST [44] v2.1.0, and ‘stats.sh’ script from BBMap [45] v38.87. Assembly contiguity was also sufficiently good with an N50 of 2,802 (see Supplementary Table S4 for additional statistics). Filtered RNA-Seq reads from all ten anemone samples were mapped to the final *de novo* assembly using Bowtie2 [46] v2.5.1, and the transcript expression levels were quantified using RSEM [47] v1.3.1.

The final *de novo* transcriptome was annotated by blasting against the UniRef90 database (downloaded on July 19, 2023; [48]) using DIAMOND [49] v2.1.8 (blastx --max-target-seqs 1 -e 1e-5). Functional annotation of transcripts with Gene Ontology (GO) terms was carried out in OmicsBox [50] v1.4.11. Additionally, eggNOG-Mapper [51] v2.1.0 with EggNOG [52] v5.0.2 was used for annotation based on orthology predictions and the results were merged with the GO annotations, resulting in a total of 21,216 (54.56%) annotated transcripts.

### Differential expression analyses

Principal component analysis (PCA) using regularized log transformed (rlog) counts was performed in R v4.2.1 to explore gene expression patterns of *A. clarkii* skin and brain samples and *E. quadricolor* tentacles and detect and remove any outlier samples. Pair-wise comparisons were carried out between the control and interaction treatments separately for skin, brain regions and tentacle samples using DESeq2 [53] v.1.32.0 in R v4.2.1 using Wald test to identify differentially expressed (DE) genes (accounting for the effect of sex for *A. clarkii*). For all pair-wise comparisons, genes with False Discovery Rate (FDR) adjusted p-value less than 0.05, absolute log 2-fold change greater than 0.3 and baseMean greater than 10 were considered to be significantly differentially expressed. Functional enrichment analyses were performed for all significantly differentially expressed gene sets using Fisher’s exact test in OmicsBox v1.4.11 with FDR corrected p-value cut-off 0.05 and the option to “reduce to most specific” GO terms.

## Results

### *A. clarkii* acclimation behaviour

Visual examination of recorded videos of fish housed with live sea anemones (n=5) revealed that the fish showed characteristic acclimation behaviours. Upon introducing the anemone into tanks, fish initially maintained distance before gradually approaching the anemone, sometimes hovering near the anemone for several minutes before making first contact. Initial contact involved fish touching anemone tentacles with their mouth or fins, often swimming away after being stung but then returning to continue acclimation. Contact frequency and intensity between the fish and anemone increased progressively as fish transitioned from tentatively touching the anemone with just their fins and mouth to rubbing their entire body against the tentacles. Additionally, fish were observed nibbling the tentacles and fanning the anemone. Within five hours of introducing the anemone into the tanks, four out of five fish were nearly fully acclimated, however one individual remained unacclimated, only touching the anemone with its caudal fin and showing avoidance. Within 24 hours of introducing the anemone to the tanks, all fish achieved full acclimation, swimming freely among the tentacles without showing any visible signs of avoidance or distress upon contact.

### Transcriptomic responses in *A. clarkii* skin upon acclimation

Upon acclimation to the anemones, significant changes in *A. clarkii* skin gene expression were observed. A total of 38 genes were significantly differentially expressed (DE) when comparing fish in control to those acclimated with anemones, with the majority (36 genes) being downregulated in fish acclimated with anemones (Supplementary Table S5). These genes were associated with diverse functional categories such as cytoskeleton, vesicle transport, metabolism, and cellular stress (Figure 2). Notably, a mannose receptor (MRC1) that binds C-type lectins and RalBP1-interacting protein 2 (REPS2), which is involved in receptor endocytosis, were both downregulated in fish acclimated with anemones (Figure 2). Functional enrichment analysis identified two significantly enriched GO terms (sarcomere and myofibril assembly) among the 38 DE genes (Supplementary Table S6).

**Figure 2:**
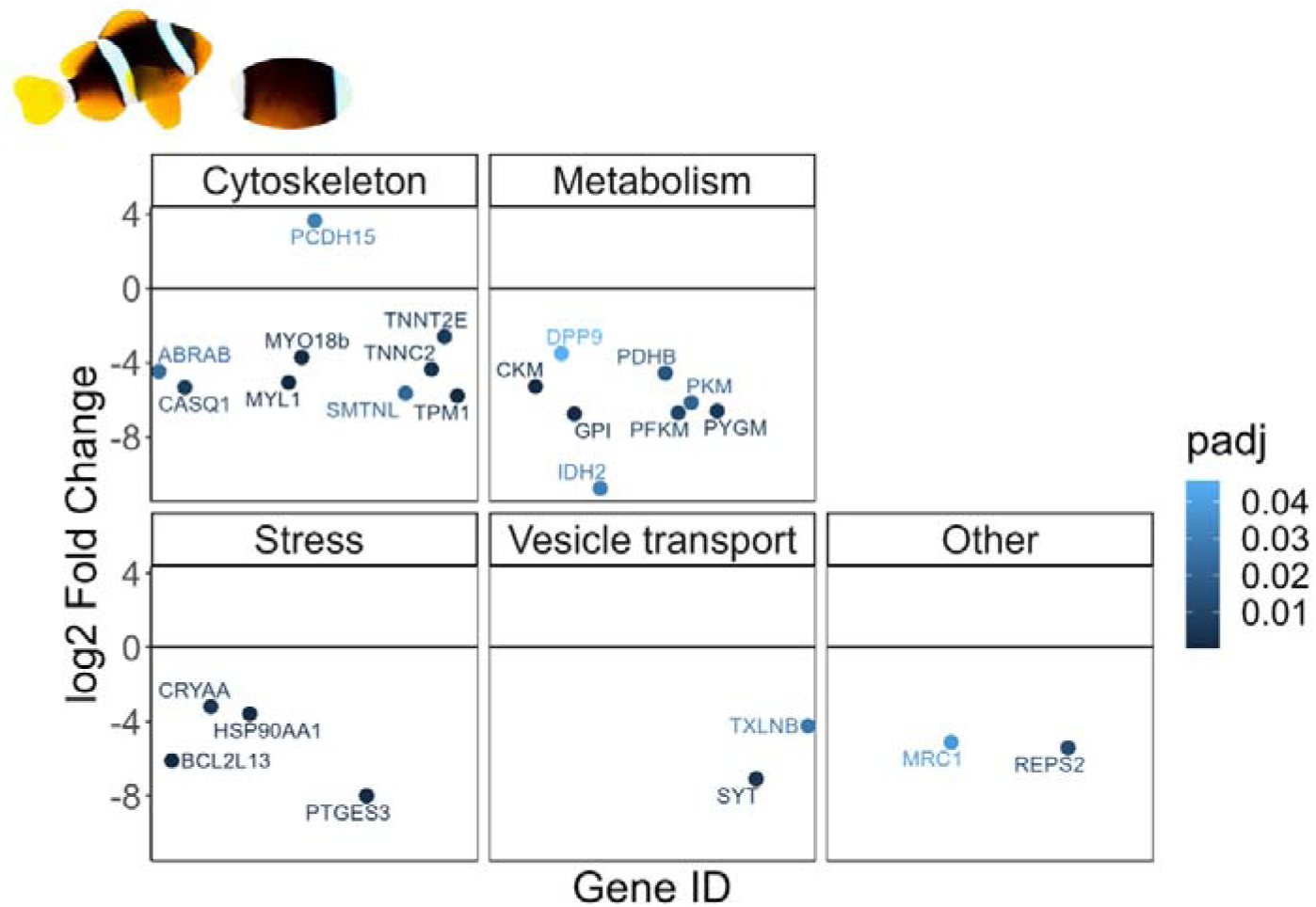
Functional categories associated with the differentially expressed genes in *A. clarkii* skin. Each point represents one gene, and the colour represents the FDR corrected p-value (padj) reported by DESeq2.

### Neurotranscriptomic responses in *A. clarkii* brain regions upon acclimation

The five brain regions (diencephalon, telencephalon, optic tectum, cerebellum, and brain stem) displayed distinct gene expression patterns upon acclimation and mutualism establishment (Figure 3(a)). The magnitude of transcriptional response varied considerably among regions, with the brain stem showing the largest transcriptional response (21 DE genes), followed by telencephalon (14 DE genes), cerebellum (9 DE genes), diencephalon (3 DE genes), and optic tectum (2 DE genes; Supplementary Table S5). Notably, there was minimal overlap in DE genes across the five brain regions (Figure 3(b)), with only one gene (PRUNE2) commonly DE among brain stem, telencephalon, and optic tectum, and one other DE gene (unannotated) shared between brain stem, cerebellum, and telencephalon. No GO terms were significantly enriched among DE genes in each individual brain region.

**Figure 3:**
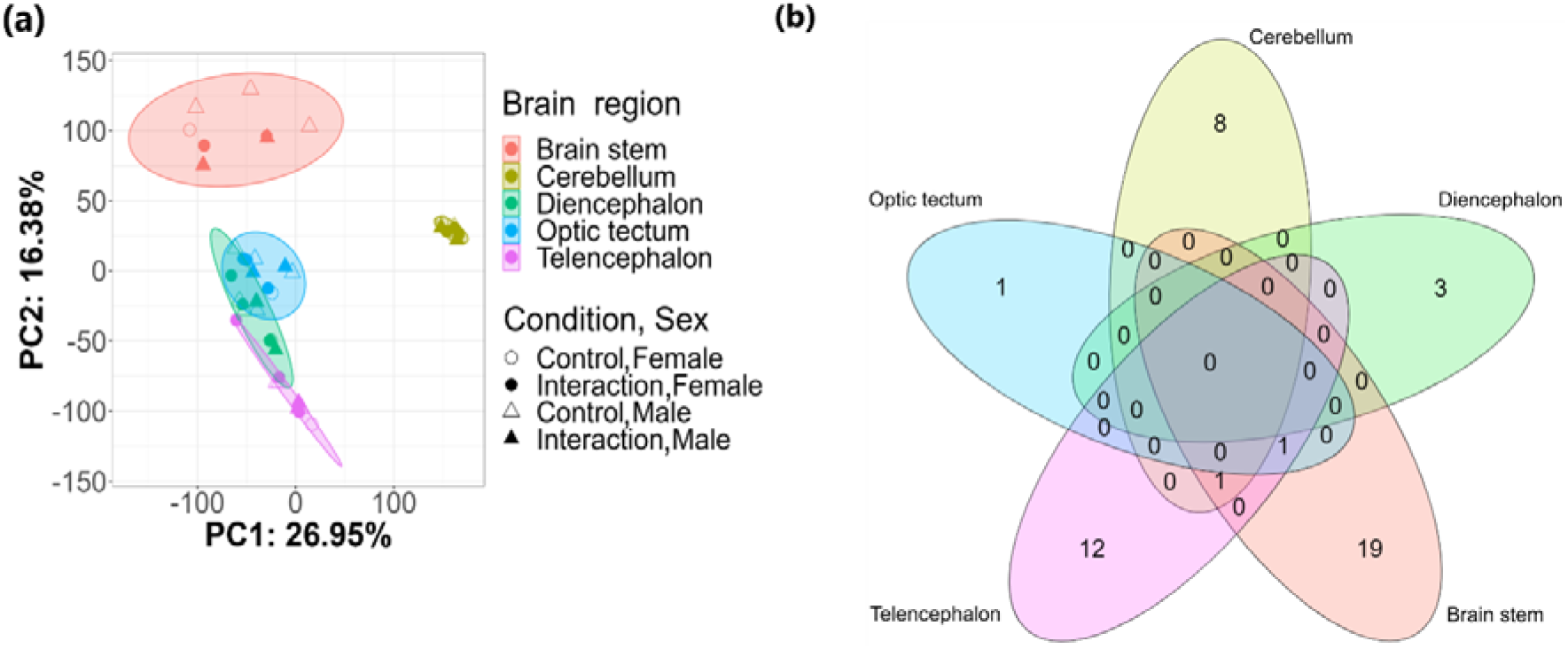
**(a)** Overall pattern of gene expression across the five brain regions of *A. clarkii* in control conditions and associated with anemones. The principal component analysis with 95% confidence ellipses was done using the regularized log-transformed (rlog) counts of all expressed genes. **(b)** Number of differentially expressed genes between *A. clarkii* in control and associated anemones for each of the five brain regions.

The forebrain regions (telencephalon and diencephalon) showed evidence of reduced cellular stress, indicated by downregulation of CYP1A, CRHBP, NRDE2 in the telencephalon, and ERCC4 in the diencephalon of anemone-acclimated fish compared to controls. Additionally, PMEL, which is involved in melanin biosynthesis and pigmentation, was upregulated in the diencephalon, and CFAP74, which is involved in olfactory signalling, was upregulated in the telencephalon of fish upon acclimation. In contrast, the hindbrain subregions (cerebellum and brain stem) exhibited differential expression of genes primarily associated with somatosensory functions, cognition, and neurogenesis (Figure 4; Supplementary Table S5). This regional specificity suggests that distinct neural functions are activated in each of the brain subregions in fish upon acclimation.

**Figure 4:**
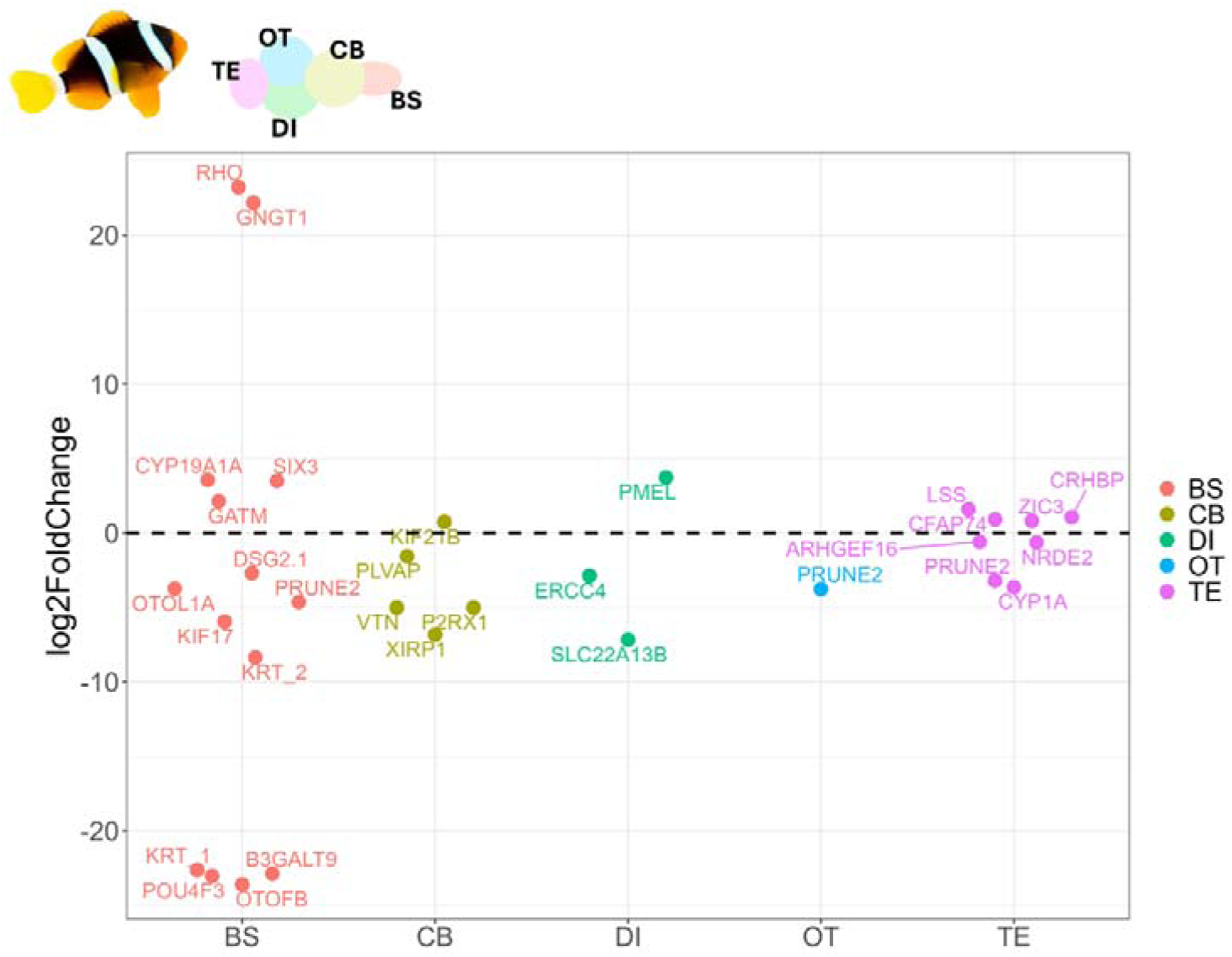
Expression changes (log2 fold change) of differentially expressed genes between *A. clarkii* in control and acclimated with anemones across the five brain regions. BS: Brain stem, CB: Cerebellum, DI: Diencephalon, OT: Optic tectum, TE: Telencephalon.

### *E. quadricolor* transcriptome profile upon mutualism establishment

*E. quadricolor* displayed a substantially more extensive transcriptional response than *A.* clarkii upon mutualism establishment, with 965 differentially expressed (DE) genes identified (578 upregulated, 387 downregulated; Supplementary Table S7). However, functional annotation was available for only 321 upregulated and 209 downregulated DE genes based on the UniRef90 database, limiting functional enrichment analysis to these annotated genes.

There were no significantly enriched GO terms among the downregulated DE genes while a total of 145 GO terms were significantly enriched among the upregulated DE genes (Supplementary Table S8). These could be grouped into eleven broader functional categories which included signalling, G-protein coupled receptor (GPCR), venom proteins, immune response, TGF-beta family, stress response, gene expression regulation, ion channels, molecular trafficking, cell cycle, and structural genes (Supplementary Figure S3; Supplementary Table S9).

## Discussion

The initial acclimation period is pivotal for establishing the mutualistic relationship between anemones and anemonefishes, during which the fish acquires protection from the host’s stinging tentacles [14,25]. In our study, all fish displayed typical acclimation behaviour upon introduction to an anemone and were fully acclimated within 24 hours. Changes in the transcriptional landscape observed in both partners upon acclimation highlight key molecular processes underlying mutualism establishment.

In anemonefish, acclimation was accompanied by modulation of sensory pathways in the brain. Differential expression of genes involved in sensory perception, signaling, and sensory memory across the telencephalon, cerebellum, and brain stem suggests that modulation of sensory processes plays a central role during acclimation. This molecular signature aligns with the known reliance of anemonefishes on olfactory cues to recognize and locate their host anemones [54–56]. Supporting this, we observed upregulation of CFAP74 in the telencephalon, a cilia- and flagella-associated protein that supports olfactory function [57,58]. Furthermore, the downregulation of XIRP1, an inhibitor of actin depolymerization, in the cerebellum may enhance associative olfactory memory in anemonefish via actin remodeling, a mechanism also observed in honeybees [59,60]. Together, these patterns suggest that olfactory learning and memory, perhaps similar to processes occurring during embryonic imprinting [54], may be reactivated in adults during reacclimation to an anemone. The detection of these olfactory cues could then trigger the behavioural responses necessary for acclimation in the fish. In fact, aromatase (CYP19A1A), which plays a role in sensory processing and sensory memory within the social behaviour network, was upregulated in the brain stem, where it facilitates translation of sensory cues to behavioural responses [61–63]. Overall, these coordinated changes in sensory genes and signaling pathways indicate that anemonefish dynamically integrate olfactory cues from their host to guide acclimation behaviour that facilitates mutualism establishment.

The most intriguing aspect of the anemone-anemonefish mutualism is the resistance of the anemonefish to anemone venom. Here we found differential expression of various cytoskeletal genes in the anemonefish skin upon acclimation, which could facilitate venom resistance by inducing structural modifications in the epidermal plasma membrane receptors potentially targeted by venom proteins [11]. This mechanism of target alteration, whereby changes in receptor protein structure reduces the binding affinity of venom proteins, represents a well-documented pathway for toxin resistance [11,64]. Cytoskeleton remodelling has also been implicated in defense response of plants against pathogens [65]. Notably, cytoskeleton genes were identified to be under positive selection at the origin of anemonefish radiation, potentially contributing to the evolution of this iconic mutualistic relationship and the subsequent adaptive radiation of anemonefishes [12]. Furthermore, other complementary resistance mechanisms were identified in the skin of acclimated anemonefish. These include downregulation of a mannose receptor (MRC1), which binds C-type lectins, a common component of anemone venom [66,67], and an endocytosis-related gene (REPS2), whose expression is correlated with receptor internalization [68]. A similar mechanism has been observed in opossums, where alterations in C-type lectin protein binding sites potentially provides protection from venomous snake predators [69]. Additionally, downregulation of synaptotagmin and taxilin, which are involved in neurotransmitter release [70–72], may provide tolerance to neurotoxins and other pore forming toxins found in anemone venom by preventing uncontrolled neurotransmitter release [67,73]. Together with previous findings that anemonefish actively reduces sialic acid levels in their mucus as a protective mechanism against nematocyst discharge [16], our results suggest that anemonefish employ multiple complementary defense mechanisms for venom resistance, thereby facilitating their mutualistic relationship with sea anemones.

Venom resistance of anemonefish is particularly crucial, as host anemones underwent substantial changes in their transcriptional landscape upon acclimation, suggesting that they mount an active response to the presence of the fish. The upregulation of genes associated with mechanosensory functions and structural components of mechanoreceptors in anemones hosting *A. clarkii* indicates activation of mechanosensory systems in response to fish movements. Anemones rely on both chemical and mechanical sensory systems to detect movements around them and discharge nematocysts in response [74], and the length and density of hair bundle mechanoreceptors in anemone tentacles increase in response to physical stimulation [75]. Furthermore, upregulation of genes associated with GPCR and hedgehog signaling, two signaling pathways known to be associated with sensory cilia in mechanoreceptors [76], further supports the activation of mechanosensory systems. This mechanosensory activation is followed by nematocyst discharge as evidenced by increased expression of genes encoding calcium and potassium channels, ion channels involved in nematocyst exocytosis [77–80], and venom proteins in anemones hosting fish. Therefore, while previous studies have proposed that anemonefish mucus molecularly mimics that of the anemone [11,17,18] and might lack certain components that triggers nematocyst firing [11,13], our findings suggest that fish movements continue to activate cnidocyte mechanoreceptors resulting in anemones actively discharging nematocysts at their mutualistic partner.

Mutualistic association induced comprehensive physiological adjustments in both species. In anemonefish, acclimation may trigger pigmentation changes, evidenced by the upregulation of PMEL in the diencephalon, a melanocyte protein involved in melanosome morphogenesis and pigmentation. While we did not directly measure phenotypic colour changes, changes in melanism is known to occur in some anemonefishes upon association with certain anemone species [9], and while their adaptive value remains unclear, it may be linked to host toxicity tolerance [81]. We also detected molecular signatures of stress alleviation in the fish following successful acclimation indicated by the upregulation of CRHBP, an inhibitor of the stress hormone corticotropin-releasing hormone, and downregulation of various other cellular stress response genes in the telencephalon, diencephalon, and skin. This stress alleviation likely reflects a shift from a state of increased predation vulnerability when fish were separated from their host anemones, similar to conditions experienced when host anemones undergo bleaching [82], to a more secure state once the mutualism was re-established. Furthermore, shifts in metabolic gene expression in the skin of acclimated fish point to metabolic changes resulting from nutrient exchange between partners [83,84]. In anemones, association boosts expression of immune response and stress response genes, likely aiding tolerance to microbiome shifts [19]. However, increased stress gene expression hints at potential hidden physiological cost for the anemone, such as oxidative stress from enhanced oxygenation and waste processing [22,84–86]. Although this might seem detrimental to the anemone, the shift in the anemone microbiome composition upon mutualistic establishment could compensate for the increase in free radical metabolites [22]. Collectively, these findings highlight that the establishment and maintenance of this mutualistic relationship involves coordinated physiological adjustments in both partners, underscoring the complexity of this mutualistic partnership.

## Conclusion

Our findings reveal that the establishment of the iconic mutualism between sea anemones and anemonefish involves coordinated molecular changes in both partners during the critical acclimation period. In *A. clarkii*, acclimation induced transcriptional shifts in genes involved in sensory perception and memory across brain regions, facilitating olfactory-mediated host recognition and acclimation. Contrary to the hypothesis that anemonefish do not trigger nematocyst discharge [11], anemones hosting fish upregulated genes encoding venom proteins and ion channels associated with nematocyst discharge, possibly due to activation of mechanoreceptors, though further research is needed to confirm the functional significance of these transcriptional changes. However, the observed differential regulation of cytoskeleton genes and venom protein receptors in the anemonefish skin may confer tolerance to the anemone venom. Both partners also exhibited molecular signatures of physiological changes which could be key in driving the mutualistic association. Overall, our results show that mutualism establishment is underpinned by dynamic transcriptomic changes in both the anemonefish and its host anemone, underscoring the complex interplay of molecular events that underlies their mutualistic association.

## Ethics

Sample collection was carried out following all institutional and national law guidelines. The experiment was completed under the approval of the Committee on the Use of Live Animals in Teaching and Research (CULATR), University of Hong Kong (# 5830-21) and the Animals (Control of Experiments) Ordinance (cap. 340), Director of Health, Government of the Hong Kong Special Administrative Region (# 21-1041/1042/1043/1044).

## Data accessibility

The RNA-Seq raw sequences are deposited in NCBI under BioProject ID PRJNA1056457. (Reviewer link: https://dataview.ncbi.nlm.nih.gov/object/PRJNA1056457?reviewer=213a93o51l1re45kcdemibqnlg). Code used for analyses is available at https://github.com/sneha100895/Anemone-anemonefish-mutualism-RNA-Seq

## Declaration of AI use

The authors used OpenAI’s GPT-5 (ChatGPT) and Claude Sonnet 4 for minor wording improvements. No unpublished or sensitive data were shared.

## Author contributions

The experiment was designed by SS and CS and run by SS with input from CS and help from DR. Anemone and fish dissections were carried out by all authors. Molecular lab work was performed by SS. SS carried out the RNA-Seq data analysis with input from CS. SS lead the writing of the manuscript with input from CS and all authors read, edited, and approved the final manuscript.

## Acknowledgements

We thank Dr. Jingliang Kang, Dr. Arthur Chung, Dr. Jade M. Sourisse, Sandra Ramirez-Calero, Helen Leung, Kam Yan Chit, and Ho Wu Cheuk for helping with the collection of the *A. clarkii* samples and Dr. Taylor A Bogar and Jiaxin Hu for providing the cameras. A special thanks goes out to Dr. José Ricardo Paula for providing valuable insights for improving fish care and experimental design. We are grateful to Kam Yan Chit and Munisa Tabarova for helping with the daily care of the fish. We also thank all members of the laboratory for their support.

