## Supplementary Figure for "Molecular drivers of mutualistic association between anemone and anemonefish"

**SUPPLEMENTARY FIGURES**

**Supplemenary Figure 1:**

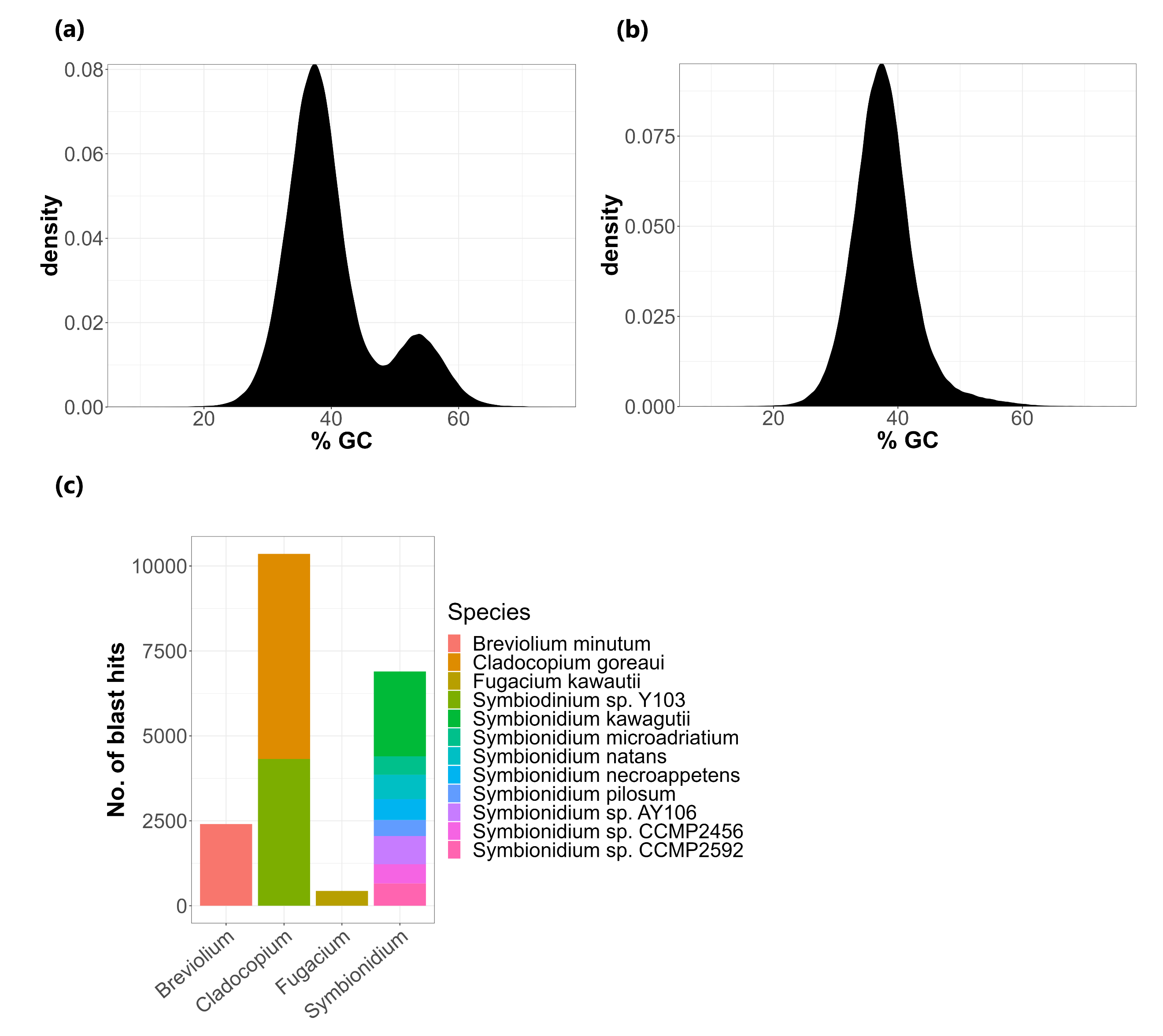

**Figure S1:** Identification and removal of Symbiodiniaceae from the *E. quadricolor* transcriptome assembly. (a) GC content of the initial transcriptome assembly. The peak with ~37% GC corresponds to *E. quadricolor* transcripts and the peak at ~53% corresponds to Symbiodiniaceae transcripts. (b) GC content of the “cleaned” transcriptome assembly after removing Symbiodiniaceae transcripts. (c) Symbiodiniaceae species identified in the *E. quadricolor* RNA-Seq data.

**Supplemenary Figure 2:**

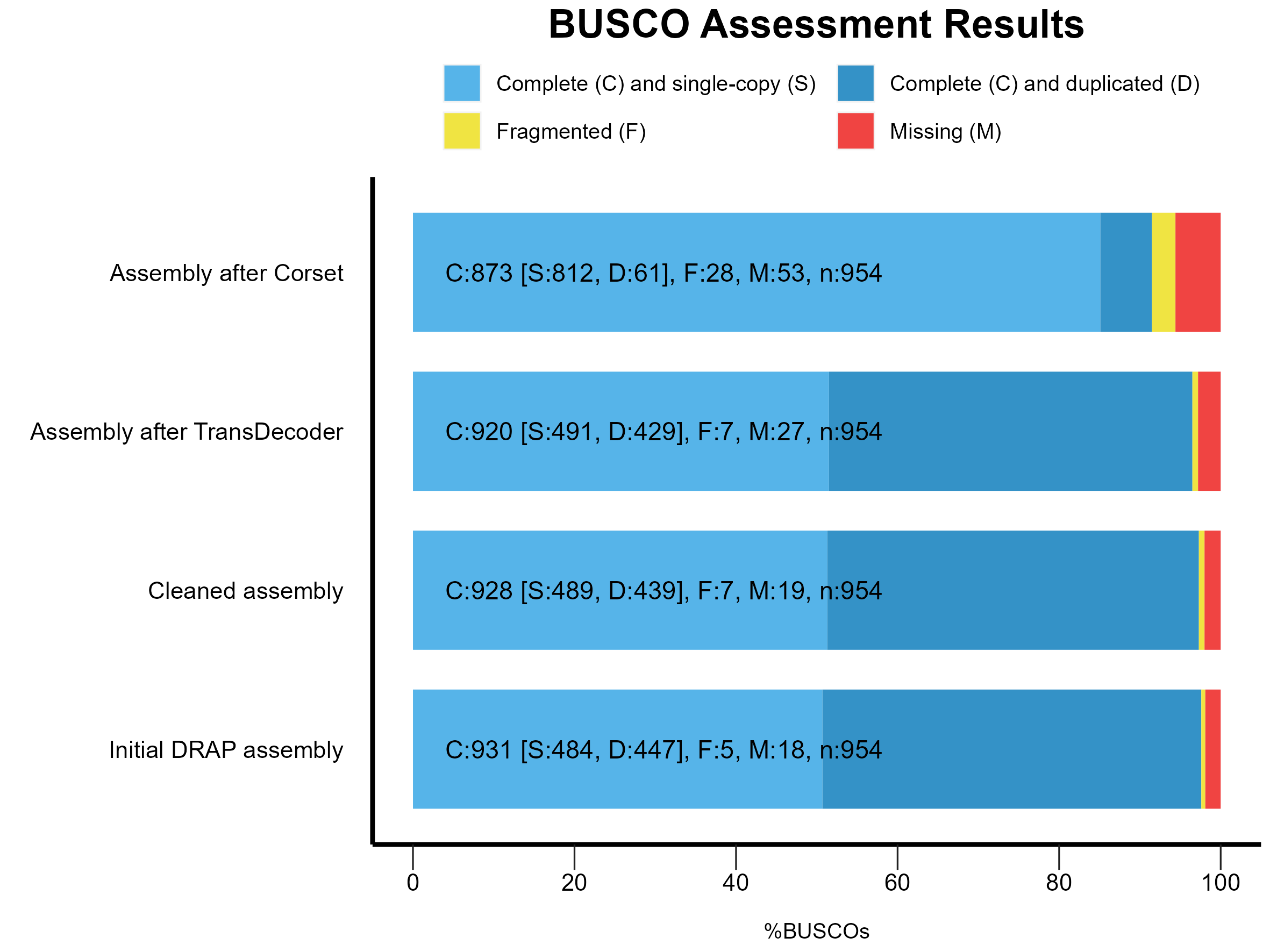

**Figure S2:** Assessment of assembly completeness using BUSCO using the Metazoa_odb10 database. The initial assembly from DRAP had a high level of completeness with > 90% of genes from Metazoa_odb10 database being recovered and this was retained in the cleaned assembly (after removing dinoflagellate transcripts) and after removing transcripts without coding potential using TransDecoder. Hierarchically clustering of transcripts using Corset greatly improved the assembly quality as seen by the reduced number of duplicate transcripts.

**Supplemenary Figure 3:**

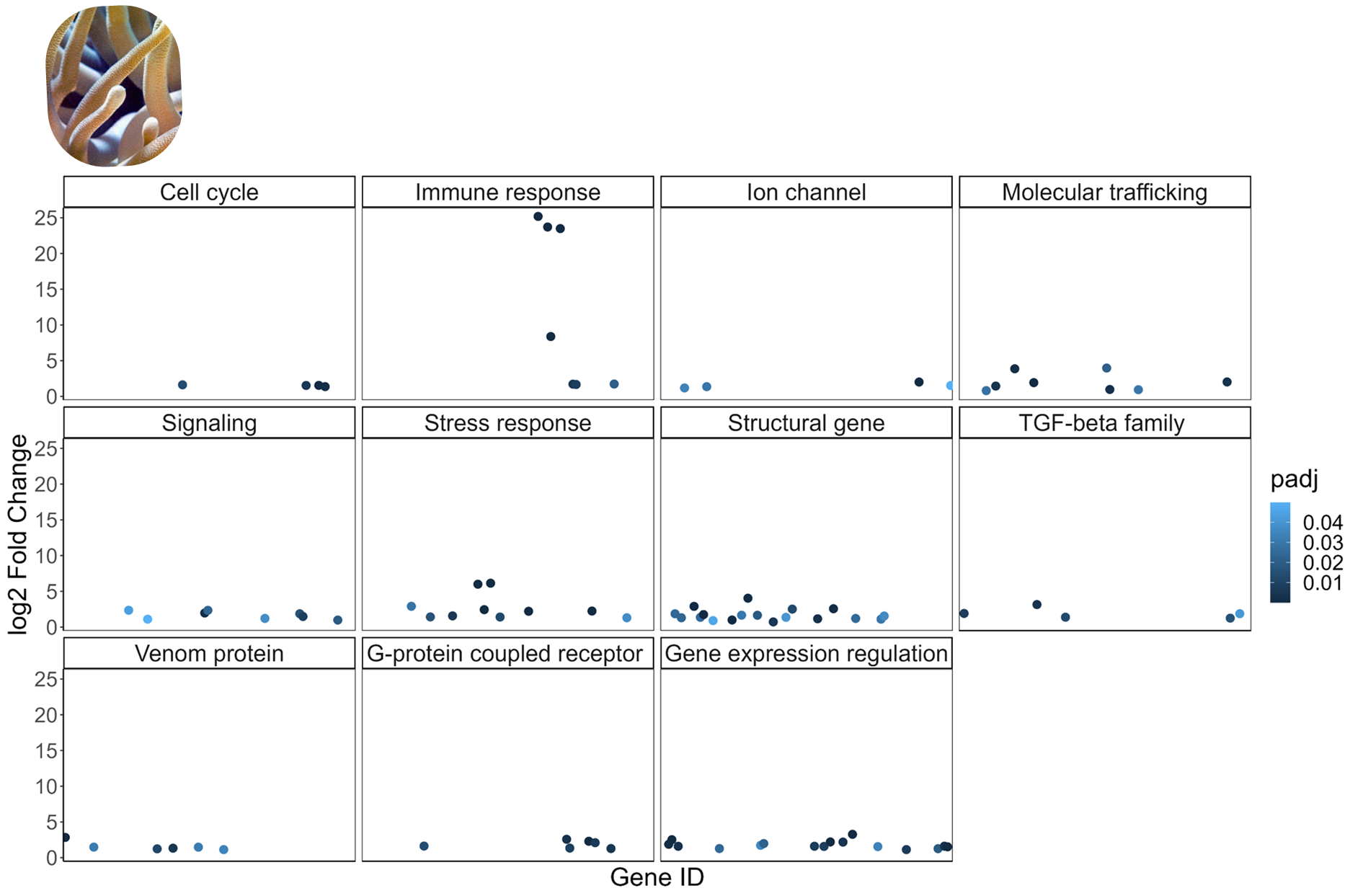

**Figure S3:** Functional categories associated with the differentially expressed genes in *E. quadricolor* tentacles. Each point represents one gene, and the colour represents the FDR corrected p-value (padj) reported by DESeq2.
